# Farming with Alternative Pollinators for Increased Biodiversity and Smallholder Incomes in Zimbabwe

**DOI:** 10.1101/2024.11.29.626070

**Authors:** Sophie Allebone-Webb, Felix Gossrau, Chloé Orland, Gracian Bara, Cédric Fioekou, Annah Matsika, Karl Riber, Bader Mahaman Dioula

**Author notes:** Corresponding author: Chloé Orland. Paper submitted to the International Journal of Agricultural Sustainability November 2024, revised version submitted 05/02/2025. Preprint version available at* https://biorxiv.org/cgi/content/short/2024.11.29.626070v1.

## Abstract

Pollinator populations have dramatically declined over the past 50 years, with over 40% of invertebrate pollinator species at risk of extinction largely due to intensive agriculture, pesticide use, habitat loss and climate change. Pollinators provide an essential ecosystem service, with about 75% of global crops relying on pollination by animals. It is therefore essential to reconsider conventional farming practices, which are largely responsible for this decline. By cultivating flowering crops “Marketable Habitat Enhancement Plants”, (MHEPs), alongside the edges of pesticide-free fields, the Farming with Alternative Pollinators (FAP) approach aims to enhance the presence of wild pollinators. In this study, we compared the performance of smallholder farmer plots using the FAP approach with plots following conventional approaches, for pollinator abundance and diversity, yield and income for 43 plots in Zimbabwe. We found significantly higher pollinator abundance and richness in FAP plots compared to control plots. There was significantly higher income and higher value of yields for all offtake in FAP plots for both crop cycles measured. Plots with higher pollinator abundance showed significantly higher income from all crops and significantly higher value of yields, showing a clear link between pollinator populations, crop production and income.

## 2. Introduction

Pollinator populations are declining globally, particularly in agricultural landscapes (Biesmeijer et al., 2006; Gill et al., 2016; Goulson, 2019; Goulson et al., 2015; Potts et al., 2010), largely due to multiple anthropogenic pressures such as land-use changes, agricultural intensification, pollution (mainly by pesticides and fertilisers), pathogens, invasive species, and climate change (Gill et al., 2016; Potts et al., 2010; Sánchez-Bayo & Wyckhuys, 2019). An estimated 87% of flowering plants are dependent on pollinators, with this figure increasing to 94% in tropical communities (Ollerton et al., 2011). Consequently, declines to pollinator populations also impact the ecosystem services provided by these flowering plants and all those that depend on them, and can lead to interlinked degradation, cascades of extinctions, poverty spirals, and eventually “Pollinator Loss Syndrome” (Christmann, 2019; Dirzo et al., 2014).

Pollinator declines are particularly evident in agricultural landscapes, and yet 75% of food crops depend, at least partly, on animal pollination for fertilization (Klein et al., 2007). Pollination services for food production are thus vital for food production, with an estimated value of €153 billion in 2005 (Gallai et al., 2009), as well as for maintaining genetic diversity and resilience (e.g. to climate change) (Christmann & Aw-Hassan, 2012), and yet agriculture itself can negatively impact the habitats and resources needed to sustain pollinators. In addition to the drivers above, agriculture and homogenous landscapes may also negatively impact pollinators through: 1) shorter flowering periods of homogenous crops that may be less than the time needed for pollinators to complete their life cycle; 2) monoculture crops that may not be suitable for specialist pollinators (those with narrow floral choices); and 3) pollinator independent crops (e.g. cereals such as wheat) which do not provide nectar or pollen for pollinators. Addressing drivers of pollinator decline and strengthening the protection and promotion of pollinator populations and pollinator diversity particularly in agricultural landscapes is therefore critical.

Farming with Alternative Pollinators (FAP) is an approach to simultaneously conserve and promote wild pollinators (i.e. ‘alternative’ pollinators to managed honeybees) and improve production and farmer incomes. Developed by the International Center for Agriculture Research in the Dry Areas (ICARDA) (Christmann & Aw-Hassan, 2012), FAP makes use of marketable habitat enhancement plants (MHEP) cultivated alongside the edges of farmer fields. The planting of MHEPs in FAP plots aims to improve pollinator habitats, thus increasing the diversity and abundance of wild pollinators, and consequently crop production both in terms of quantity and quality. The goal is that the resultant, visible increases in farmer income per surface area will motivate farmers to permanently adopt pollinator-friendly farming practices.

The FAP approach builds on evidence showing that native wild flowering plants are important for the conservation of pollinator populations in farmlands (Dicks et al., 2015; Nicholls & Altieri, 2013) and that sown wildflower strips also attract and provide habitat for wild pollinators, increasing the abundance and diversity of bumblebees (Carvell et al., 2007). Over the last two decades, wildflower strips along fields have been introduced in several European countries within the Agri-Environmental Schemes (AES) framework, with positive impacts to biodiversity and ecosystem services (Haaland et al., 2011). However, the high and continued implementation costs of these schemes have led to doubts over their broader accessibility (Batáry et al., 2015; Christmann, 2020; Uyttenbroeck et al., 2016), particularly for low and middle-income countries. The FAP approach uses MHEPs to provide similar resources to pollinators as wildflower strips, but with additional focus on benefits to farmer incomes and improving farmer motivation (Christmann et al., 2022). Key elements in FAP approaches are crop diversification through a combination of pollinator-friendly main crop and MHEPs, temporally sequencing the planting of crops to achieve flowering overlap, habitat enhancement for pollinators by reducing harmful practices (e.g. pesticides, insecticides) and conservation agriculture practices.

ICARDA trials in Morocco and Uzbekistan have demonstrated a visibly higher yield and higher income of FAP fields compared with control plots, alongside higher pollinator abundance and diversity in FAP plots (Bencharki et al., 2023, 2023; Christmann, 2017; Sentil, Lhomme, et al., 2022; Sentil, Reverté, et al., 2022). The visible socioeconomic benefits through FAP consequently motivated farmers to adopt the approach and conserve pollinators. Inspired by this research, Action against Hunger (ACF)^4^ initiated a FAP research project in Zimbabwe to validate the approach in a small-holder context in sub-Saharan Africa.

The agricultural production system in Zimbabwe is dominated by the high use of agrochemicals (herbicides, fungicides and insecticides, unsustainable monoculture cropping), tree cutting for firewood and tobacco curing, and agricultural expansion, requiring clearing of land (e.g. deforestation, bushfires) (Mudimu et al., n.d.; Zimba & Zimudzi, 2016). These practices all have a negative impact on the wild pollinators which provide the majority of pollination services (Chakuya et al., 2022). This threatens biodiversity and adversely affects already resource-poor smallholder farmers who rely on ecosystem services such as pollination for their agricultural production, food and nutrition security and economic livelihoods. Therefore, to address the negative impacts of the practices of conventional farming on biodiversity and particularly on pollinator loss, there is a need to develop and promote alternative and sustainable solutions to conventional farming practices. The FAP method was selected (in conjunction with the use of other agroecology and sustainable agriculture techniques) for its holistic approach to farming which includes both socioeconomic and ecological benefits, and because the method itself was deemed in conjunction with local farming techniques.

This paper presents the results of FAP trials in two districts in Zimbabwe, Gokwe North and Gokwe South. The objective was to test the replicability and impact of the FAP approach in Zimbabwe. We hypothesised that using the FAP method to improve pollinator habitat would have a positive impact on pollinator abundance and diversity, and consequently have tangible benefits on crop yield (quantity and quality) and income of smallholder farmers in Zimbabwe. This research effort aims to produce scientific evidence to contribute to the existing FAP knowledge base as generated from other contexts. To our knowledge it is the first such study from sub-Saharan Africa.

## 3. Materials and methods

### 3.1. Site description

The baseline study was conducted in two districts: Gokwe North and Gokwe South, Midlands Province, Zimbabwe (Figure 1). Gokwe North lies north of Midlands Province and is in Natural Ecological Region IV. It experiences arid to semi-arid conditions, receiving between 250 and 800mm of rainfall annually. Gokwe South lies North-West of Midlands Province, with 40% of the district falling under Agro-ecological Region IV while 60% is in Region III, which is characterised by low and erratic rainfall patterns.

**Figure 1.**
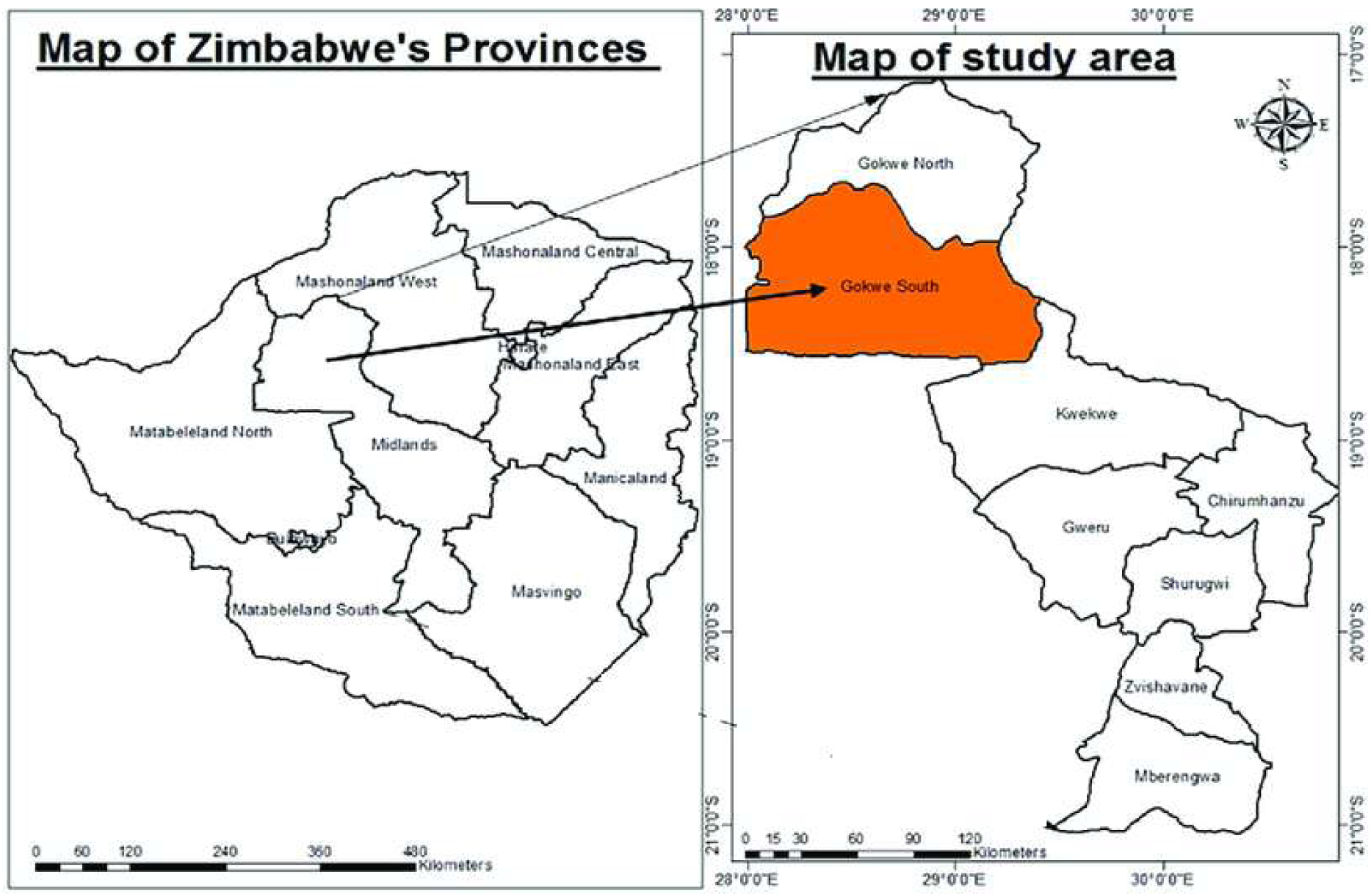
Map of the study area Gokwe North and South district. Adapted from Philips (2018), Cartographic Unit, the University of the Witwatersrand. South Africa.

### 3.2. Participants and training

The project registered and trained 80 (46 male, 34 female) project participants, with 40 participants per district for the two districts (Gokwe South and Gokwe North, Zimbabwe). Garden materials were distributed to the 80 farmers, consisting of diamond mesh wire for fencing the garden, treated poles and nails for fencing, a hoe, a watering can, and vegetable seeds. Control farmers received seeds for the main crop only, whilst FAP farmers received seeds for the main crop and MHEPs (8 different species: mustard rape (*Brassica juncea*), green pepper (*Capsicum annuum*), tomatoes (*Solanum lycopersicum*), cucumber (*Cucumis Sativus*), okra (*Abelmoschus esculentus*), pumpkin (*Cucurbita* spp.), butternut (*Cucurbita moschata*), coriander (*Coriandrum sativum*) and watermelon (*Citrullus lanatus*). See Table 1. Cropping cycles). MHEP crops were selected based on attractiveness to pollinators, farmer suggestions, and flowering periods, to ensure that MHEP crop flowering partially overlapped with main crop flowering period.

**Table 1.** Cropping cycles.

|  | Cycle 1 | Cycle 2 | Cycle 3 |
| --- | --- | --- | --- |
| Date | Sep 2022 – Feb 2023 | Mar-Jun 2023 | Jul-Oct 2023 |
| Pollinator data collection | January/February 2023 | End of June 2023 | Beginning October 2023 |
| Main crop(s) | Tomato<br>Mustard Rape | Okra | Tomato<br>Watermelon |
| MHEPs | Watermelon<br>Butternut<br>Cucumbers<br>Green Pepper | Green pepper<br>Cucumber<br>Sugar bean<br>Mustard rape<sup>8</sup> | Green pepper<br>Cucumber<br>Sugar bean<br>Mustard rape |

<sup>8</sup> One person also grew tomato in cycle 2

All participating farmers received training on the basics of the FAP concept, biodiversity, pollinator identification and counting, and field designs in FAP, as well as conservation agriculture (CA) practices for planting times, cropping density, and other aspects. The project was set up so that all farmers participating in the study used similar practices and all had access to water for irrigation. Similarly, farmers were asked not to use any fertilisers, pesticides or other amendments. However, data was not collected on actual planting densities used, and it is unlikely that pesticide or fertiliser use would have been reported, had it been used. Specific data on irrigation was also not collected.

Training was conducted by Environment Africa (EA) and Nutrition Action Zimbabwe (NAZ) who were partners of Action against Hunger in the FAP trials. Project staff and local agriculture extension agents were also trained in the FAP approach and pollinator identification and counting. A qualitative assessment of farmer perceptions towards this novel practice was also carried out to assess the buy-in from local communities (Annexe 3).

### 3.3. Design of the participatory fields

As per the ICARDA trials, in FAP plots, farmers participating in this research project allocated 25% of their plot to cultivating MHEPs, while the remaining 75% of the plot was cultivated with a pollinator-dependent main crop. This is compared with control plots where the main crop was cultivated over 100% of the plot area (Figure 2). All plots were set up to be the same size (30 x 10m), which is similar to the average size of small-holder plots in Zimbabwe and in Sub-Saharan Africa in general where most plots are less than 0.5 hectares. Each FAP plots had a 1m border of MHEP plants on each of its four edges. Control plots were the same size, but with only one crop (“main crop”) cultivated. All selected plots were a minimum of 2km apart from each other, to avoid plot conditions impacting neighbouring plots. This corresponds with pollinator behaviour: most wild pollinators generally work in an area of approximately 50 – 2000m radius from the nest.

**Figure 2.**
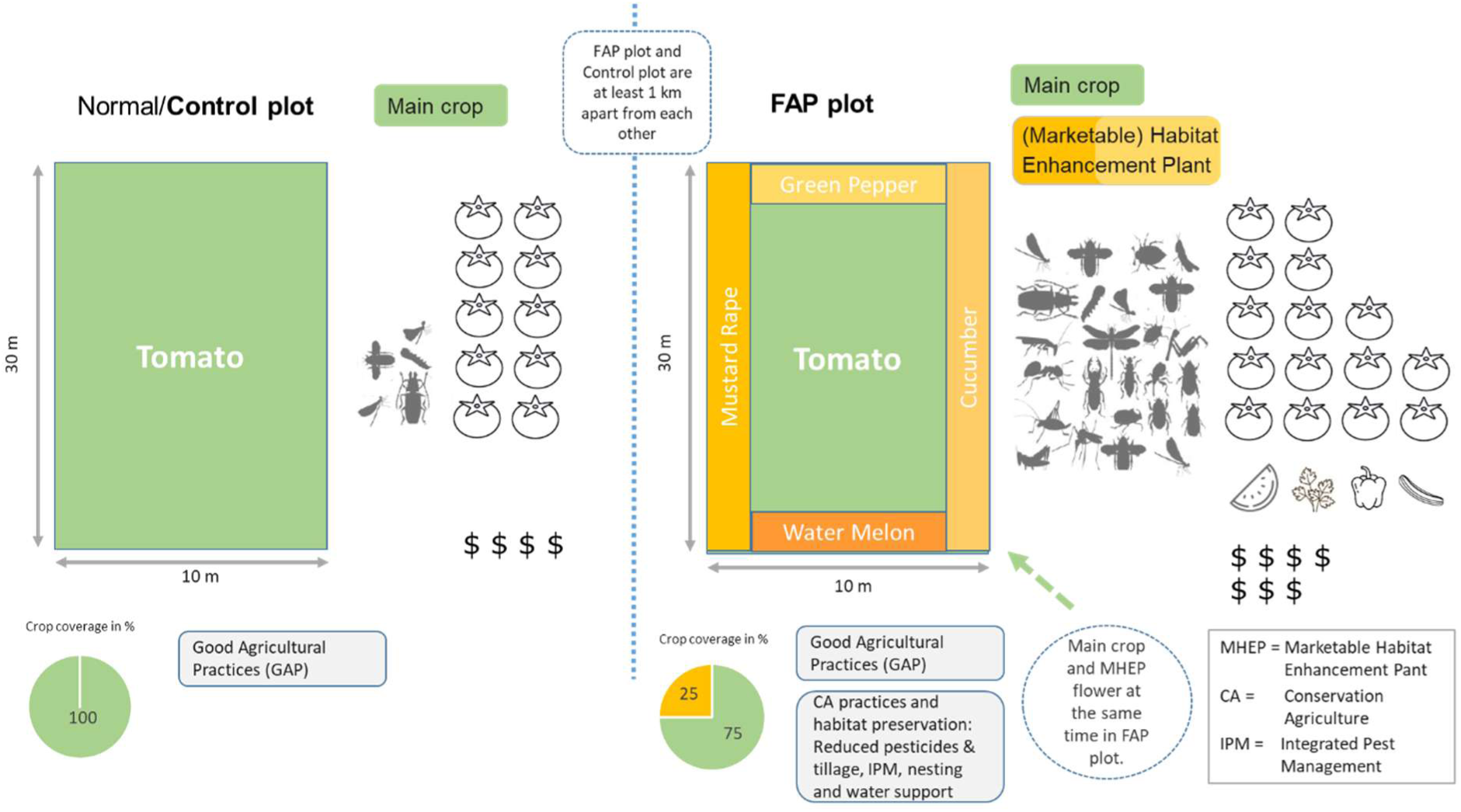
Principles and expectations using FAP demonstration and control plots

### 3.4. Cropping cycles

Table 1Data from three cropping cycles were collected from September 2022 to October 2023. Main crops and MHEPs varied depending on the cycle (Table 1). FAP farmers also included a small strip of coriander as habitat enhancement plant (but yield or income was not measured for coriander). The first cropping cycle began in September 2022, and all farmers managed to plant the five different crops in their 30m by 10m gardens. However, in January 2023, cyclone Freddy caused widespread damage in both districts, with over 50% of farmers reporting extensive damage to crops, and the rest also reporting mild or low damage. Consequently, data from the first cycle were not included in analyses on yield and income. Data from cycle 1 were included in the pollinator counts, as these were conducted before the cyclone. As such, there were 3 cycles for the pollinator data, but only the 2^nd^ and 3^rd^ cycle for income and yield data.

### 3.5. Sample size

An original 80 plots were included in this pilot study, 50 FAP and 30 control plots. Of these 80 plots, the following were excluded from the analysis:

- All community plots were removed from the analysis (nearly half) because many were not set up following the correct experimental design, or did not follow the recommended farming techniques.
- As described previously, data from cycle 1 were not included in analyses of impacts on yield or income.
- In addition, there were no data on yields or income for 14 plots points (from 8 farms), so these were also removed, leaving 117 data points from 43 farms.
- For analyses on the value of the yield (of all crops, main crop or MHEPs), only limited data were available, so there are fewer data points than for main crop yield or total income.

### 3.6. Measures of pollinators

For each plot, three timed transects were conducted. Pollinator monitors walked the 1m wide transects (Figure 3) the length of the plots over 7.5 minutes counting and recording all pollinators that landed on flowers for at least 0.5 seconds. Full procedure guidelines are shown in Annexe 1.

**Figure 3.**
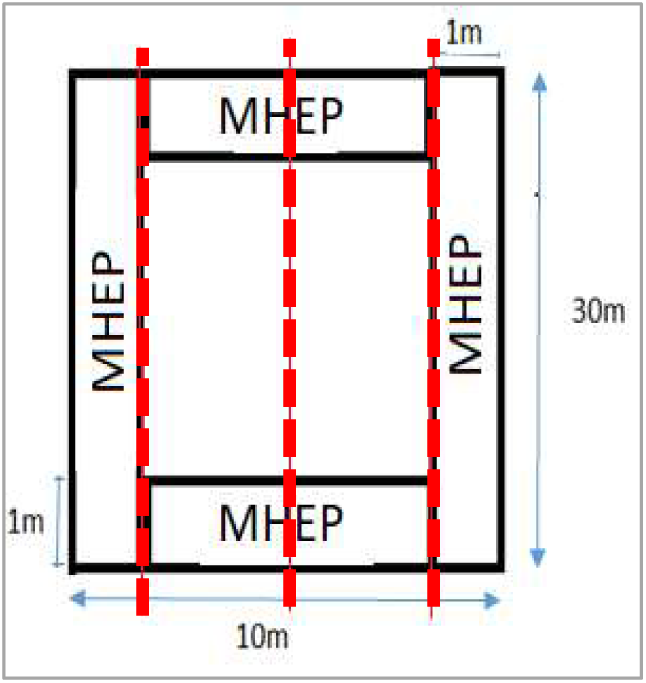
FAP plot with pollinator transects shown in red. Transects for control plots were in the same positions.

Pollinators were identified to family, genus and species level where possible using dichotomous keys by Eymann *et al* (2010) and Goulet & Huber (1993). Pollinator diversity was calculated using the Shannon-Weiner diversity index while the pollinator abundance was determined using pollinator counts.

Three measures of pollinators were included in the analysis:

- **Pollinator abundance:** total count of pollinators for that plot in that cycle
- **Taxonomic richness:** number of different taxa (identified to the lowest possible level) recorded for that plot in that cycle
- **Pollinator diversity:** pollinator abundance and richness were used to characterise pollinator diversity and account for evenness across samples using the Shannon-Weiner diversity index (H). The higher the value of H, the higher the taxonomic diversity within a particular community.

### 3.7. Yield and income assessments

Farmer reports of total yield for each crop were recorded and so may include inherent farmer recall inaccuracies. Yield was reported in Kgs harvested for all crops except mustard rape which were measured in number of bundles harvested. Farmer recall of the amount of each crop consumed or sold (in Kg or bundles) was also recorded. Similarly, the amount sold (in Kg or bundles) and the income (in USD) for each crop was recorded. Three different measures were analysed:

- **Yield:** the number of kgs or bundles recorded as produced per crop, per plot, per cycle.
- **Yield value**: in order to compare yield across crop types (i.e. for plots with multiple crop plants, and across plots with different crops), the total amount of each crop produced was taken and multiplied by the average price per unit of that crop. Total value of yield for all crops from a particular plot would therefore be the monetary value (in USD) of the main crop plus the value of all MHEP crops produced from that plot (regardless of how much was actually sold). Average prices for each crop were calculated from the data recorded across both districts and both cycles.
- **Income:** the amount of income gained (in USD) for that particular crop and plot. This does not include the value of any produce that was not sold (e.g. used for own consumption).

### 3.8. Data analysis

Statistical analyses were carried out using R 4.4.1 (R Core Team, 2024). The packages stats (R Core Team, 2024), ggpubr (Kassambara, 2023), ggplot2 (Wickham, 2016), correlation (Makowski et al., 2022), pspearman (Savicky, 2022) were used.

### 3.9. Ethical considerations

<u>Statement on ethical approval:</u> The authors confirm that the FAP project in Zimbabwe was ethically approved by ACF-France (*Action contre la Faim*), and as with all ACF actions, followed strict ethical guidelines, and conformed to the framework of fundamental humanitarian principles as enshrined in the Fundamental Humanitarian Standard (CHS). In addition, the project also followed the ACF Ethics and Research Principles and Guidelines^5^ (ACF International, 2014), and the ACF Data Protection Policy^6^, (ACF-France, 2024). This study was funded by a Darwin Initiative Innovation under grant DARNV007, and thus also conformed to the key principles of good ethical practices, including the use of “Prior Informed Consent (PIC) principles with communities”. All data was anonymised.

<u>Statement on informed consent:</u> The authors confirm that in accordance with ACF and Darwin policy, all participants gave fully informed and free verbal consent to participate in the study and for the anonymised results of this study to be used for research and publication purposes for this article and all other future publications. Obtaining consent is part of the Survey Checklist^7^ used by the survey teams (ACF-France, n.d.).

## 4. Results

### 4.1. Impact of treatment (FAP vs control) on pollinators

In total, 43 plots were included in the analyses, of which 29 were FAP plots and 14 were control plots. After removing non usable data, data were analysed from 32 plots over all three cycles (22 FAP and 10 control plots); 10 plots over two cycles (6 FAP and 4 control plots), and only 1 plot (FAP) with data from only one cycle. Data were analysed separately by cycle, and then together for all three cycles.

Groups of pollinators belonging to 5 orders were recorded: Diptera (2), Hymenoptera (7), Lepidoptera (2), Coleoptera (1), with the Hymenoptera order having the highest number of species. The following 12 groups of pollinators were identified:

- Mining bee, sub-family *Andrenidae*
- Leafcutter bee, *Megachile spp*.
- Carpenter bee, *Xylocopa spp*.
- Butterfly, order *Lepidoptera*
- Stingless bee, *Meliponini spp*.
- Sweat bee, *Halictus spp*.
- Hover fly, family *Syrphidae*
- Honeybee, *Apis mellifera*
- Wasp, family *Vespidae*
- Fly, order *Diptera*
- Moth, order *Lepidoptera*
- Beetle, order *Coloeoptera*

For all three cycles, FAP plots had a significantly higher pollinator abundance, significantly higher taxonomic richness (number of taxonomic groups counted), and significantly higher taxonomic diversity (as measured by the Shannon Index), Figure 4. For all 9 analyses, data were analysed in the R “stats” package using a Mann-Whitney-Wilcoxon test because data were not normally distributed. Cycle 1 (n=41): pollinator abundance, W=0, p<0.001; taxonomic richness, W=21.5, p<0.001; pollinator diversity, W=48, p<0.001. Cycle 2 (n=41): pollinator abundance, W=0, p<0.001; taxonomic richness, W=84, p<0.001; pollinator diversity, W=36, p<0.001. Cycle 3 (n=35): pollinator abundance, W=0, p<0.001; taxonomic richness, W=38.5, p<0.001; pollinator diversity, W=9, p<0.001). See also Annexe 2.

**Figure 4.**
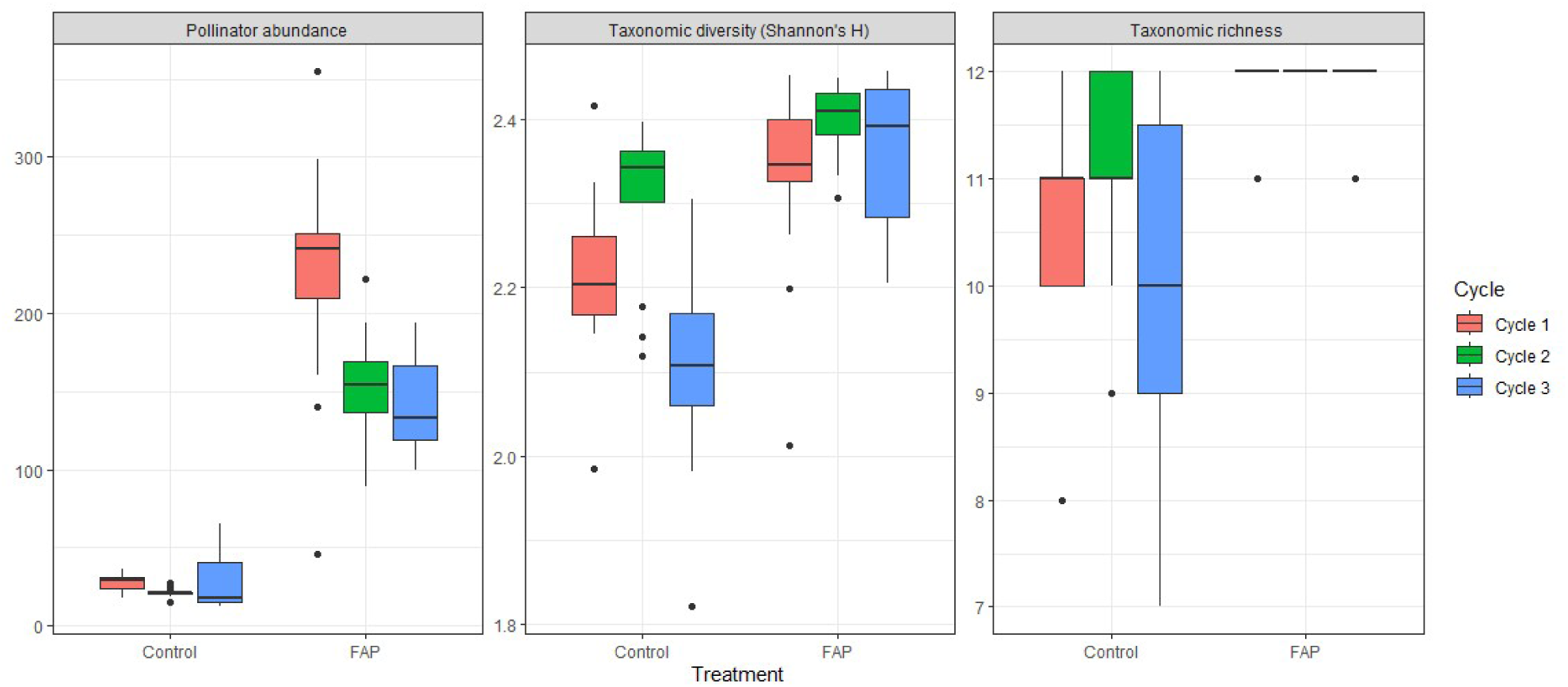
Data on pollinator abundance (a), taxonomic diversity (b) and taxonomic richness (c) for FAP and control plots for cycles 1, 2 and 3

There was a significant difference in pollinator abundance between cycles, with a higher pollinator abundance for cycle 1 compared to cycles 2 and 3 (p<0.001, n=117, DF=2). However, given the difference in crops planted and seasonal climate differences, it is difficult to know what factors had an impact on these differences. The data showed no significant difference between districts within cycles except for the 3^rd^ cycle where there were significantly higher counts of pollinator abundance for Gokwe South than Gokwe North for both FAP and control plots.

### 4.2. Impact of FAP on yield and income

#### 4.2.1. Main crop yield and income, cycle 2

To avoid comparing yields as measured by weight across different crops, comparisons of yield by plot treatment were done separately by cycle and main crop. For cycle 2, the main crop was okra for all plots. Average yield (kg) and income (USD) for okra per plot were on average higher for FAP plots, but this was not statistically significant (Mann-Whitney-Wilcoxon: yield, p=0.6639; income, p=0.30832). Given that for FAP plots, only 75% of the plot is used for cultivating the main crop (roughly 22.5m^2^), we also considered the yield and income per m^2^ for okra production. However, although FAP plots had on average a higher yield and income per m^2^, but this was also not statistically significant (yield: p=0.3063; income: p=0.2226; Figure 5). Main crop yields and income in cycle 2 were also significantly higher Gokwe North compared to Gokwe South (Yield per plot: W=352.5, p=0.000183; Income per plot: W=334.5, p=0.001069, Figure 5).

**Figure 5.**
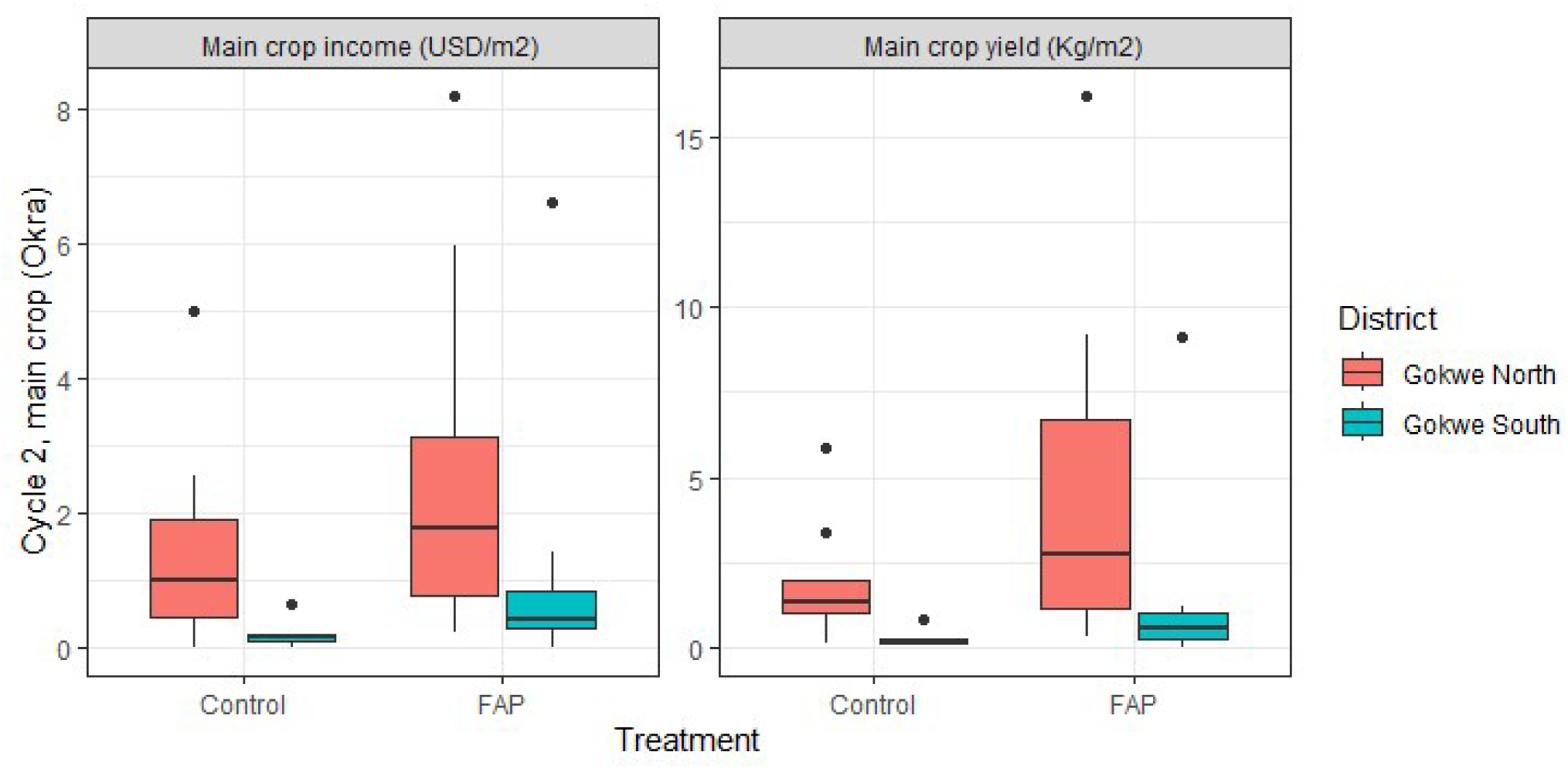
Main crop production and income per m2 for cycle 2 (Okra)

In cycle 3, farmers in Gokwe North planted tomatoes as their main crop, whereas farmers in Gokwe South planted watermelon as their main crop. Consequently, we did not analyse the differences in main crop yield or income within cycle 3, as it was difficult to distinguish the impacts of plot treatment and type of crop (different weights and prices of produce).

#### 4.2.2. Yield value and total income (main crop + MHEP crop), cycle 2

In Gokwe North, the average income/plot was higher in FAP plots: $144 compared to $43 in control plots. This was also observed in Gokwe South, where the average income was $92 in FAP plots compared to $7 in control plots. This means that the total income was 3.3 and 13 times higher in FAP plots for Gokwe North and South respectively, although some of this difference could also be explained by differences in market access (prices and amounts sold). Total income from all crops (main crop + MHEPs) was significantly higher for FAP plots in cycle 2, even when controlling for district (H=14.8, n=41, p<0.005, Figure 6). Income from all crops was also significantly higher for Gokwe North compared to Gokwe South (H=5.97, p<0.05).

**Figure 6.**
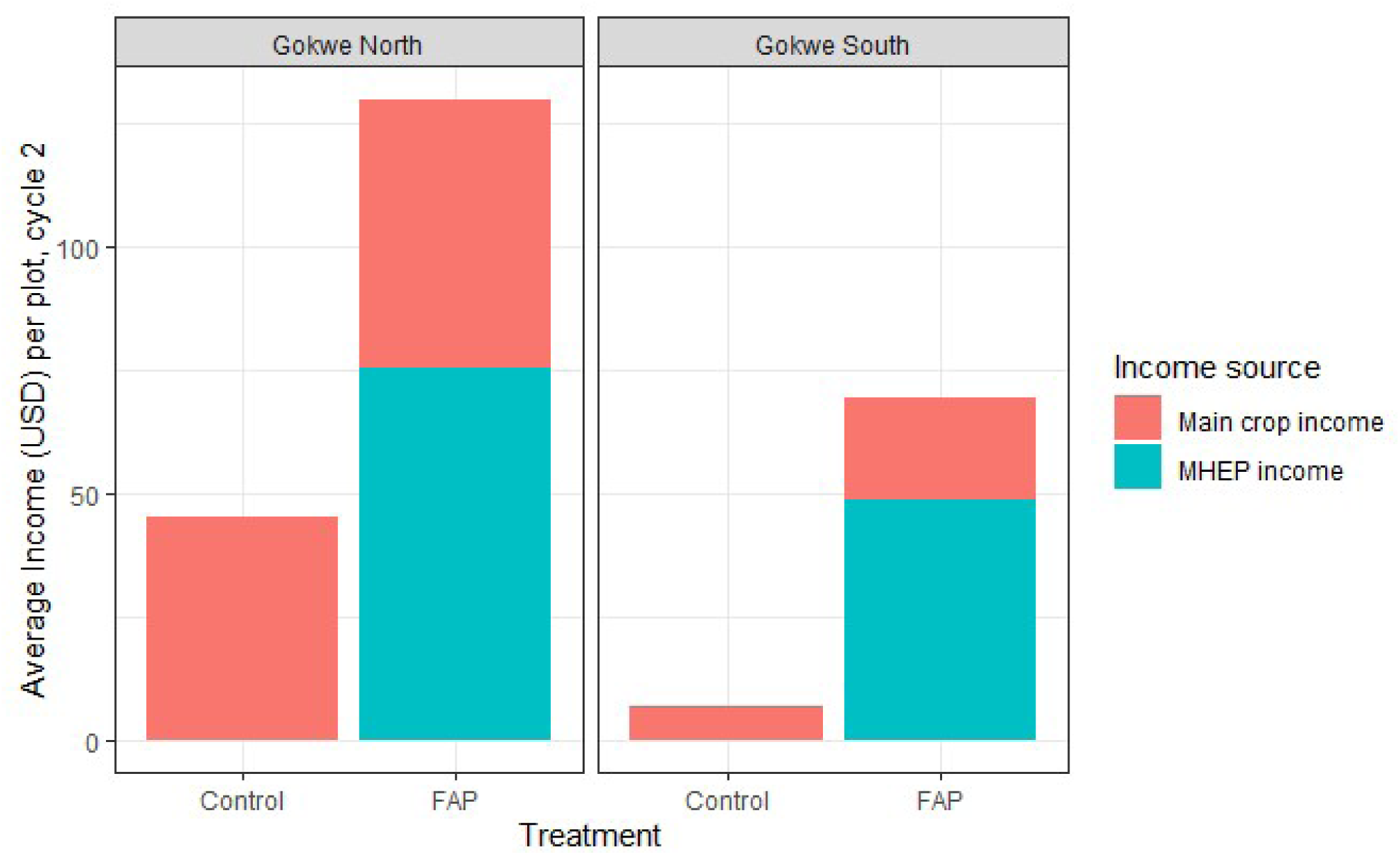
Income per plot for cycle 2 by treatment and district

#### 4.2.3. Impact of FAP on yield, yield value, and income across cycles

When controlling for main crop type, district and cycle, main crop yields are on average higher in FAP plots, but this is not significant (t=1.953, p=0.0548, n=76). However, there were significantly higher value yields from FAP plots compared to control plots, controlling for district, cycle and main crop (t=2.426, p=0.020, n=44, Figure 7).

**Figure 7.**
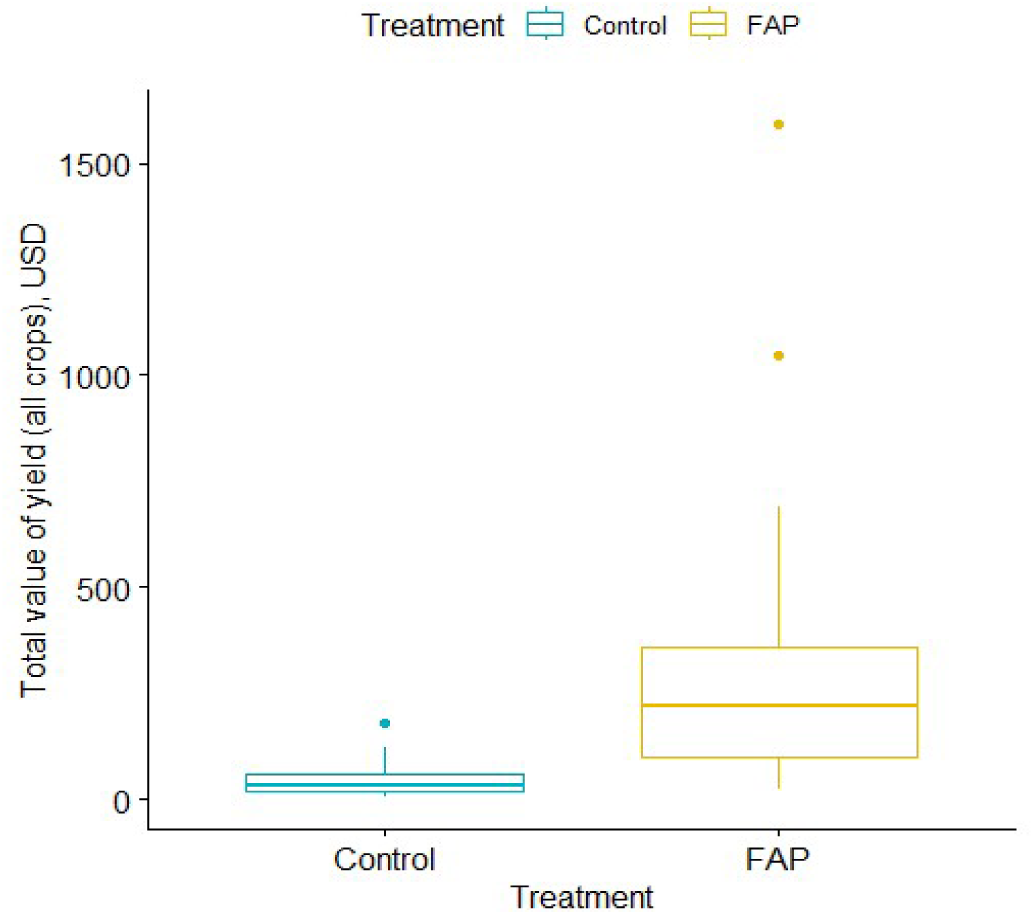
Difference in value of total yield (all crops) between FAP and control plots for cycles 2 and 3

There was a significantly higher income from all crops recorded from FAP plots compared to control plots (t=3.702, p<0.0005, n=76), again, controlling for district, cycle and main crop. Average income from main crops was higher from FAP plots compared to control plots, but this wasn’t statistically significant (t=2.002, p=0.055, n=44). These results suggest that MEHP crops provide a significant increase in income.

### 4.3. Impact of pollinators on yield and income

Pollinator abundance was positively correlated with total income (Spearmann’s correlation: rho=0.488, n=76, p<0.0001, Figure 8) and value of total yield (rho=0.628, n=44, p<0.0001, Figure 9). There were two outlier FAP plots that outperformed all other plots in both production and income. These plots were notably well maintained by very motivated farmers and were often visited as the “model” plots.

**Figure 8.**
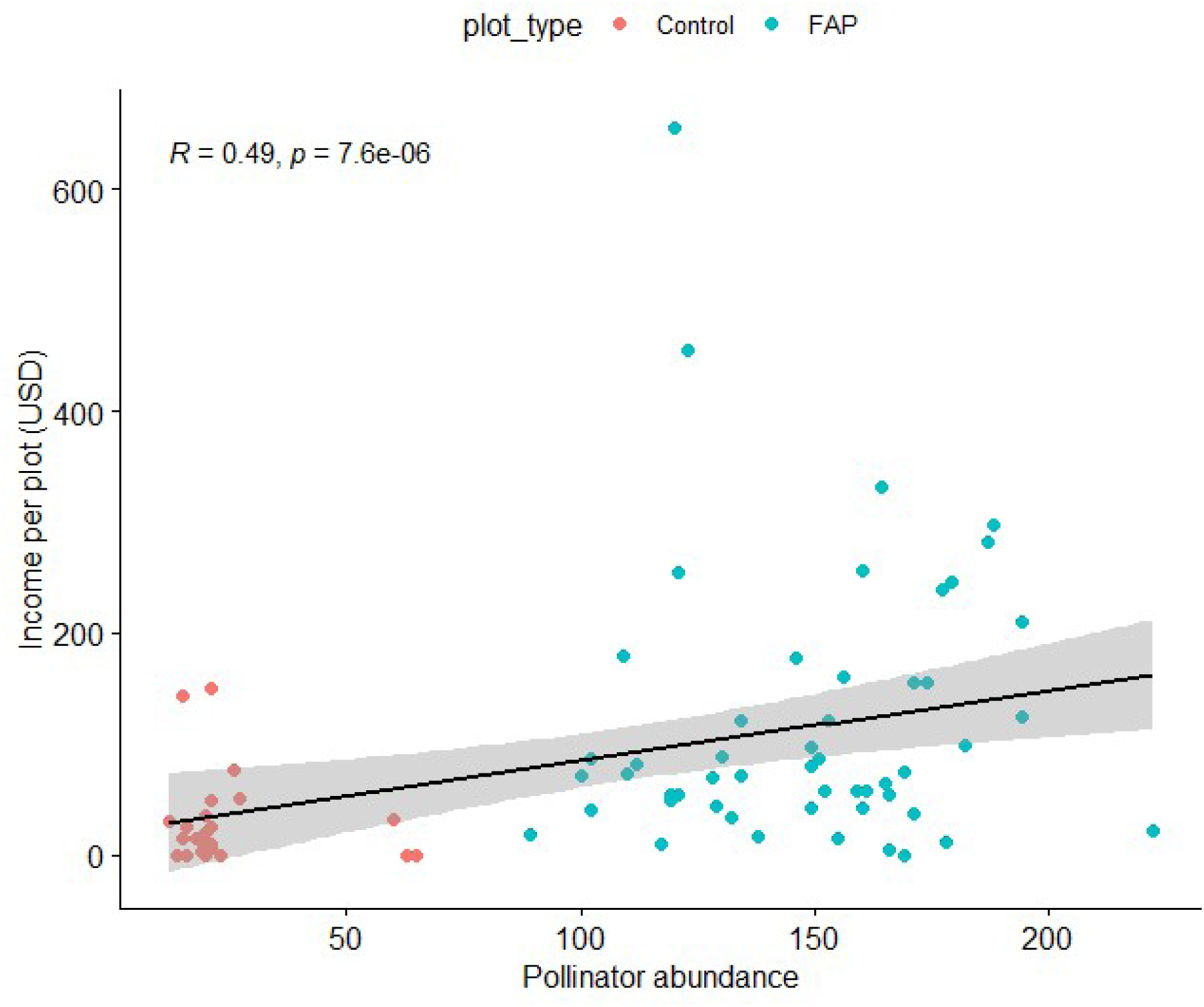
Pollinator abundance and income from all crops (main crop and MHEP) for cycles 2 and 3

**Figure 9.**
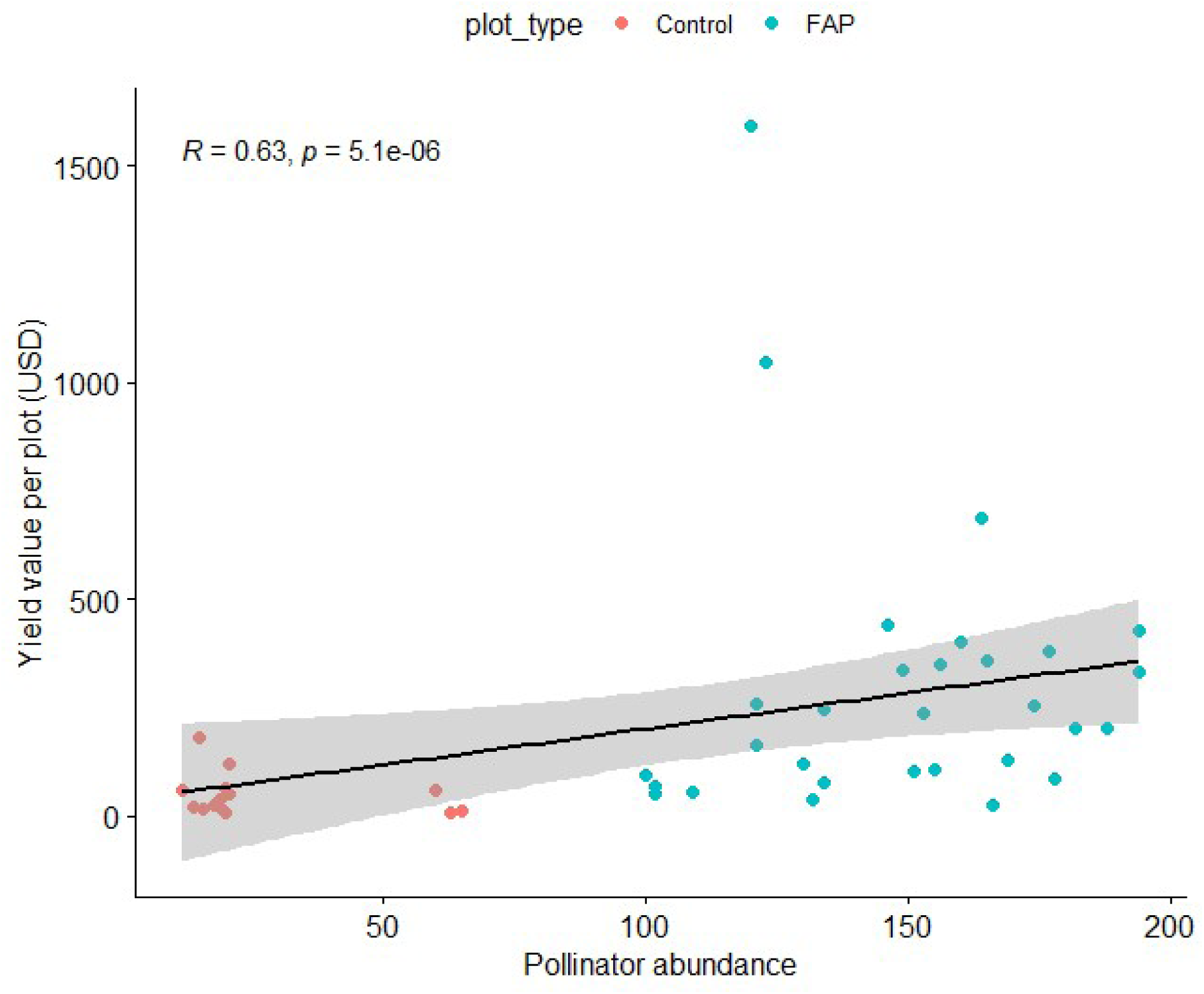
Pollinator abundance and total value of yield (all crops), for cycles 2 and 3

## 5. Discussion

In this pilot study, we tested whether incorporating pollinator habitat enhancement through MHEPs and adherence to conservation practices in smallholder agriculture plots in Gokwe North and Gokwe South, two districts of the Midlands Province of Zimbabwe, would significantly increase pollinator diversity and have impacts on crop yields and income. Our results are encouraging, with significantly higher counts and diversity of pollinators seen in FAP plots compared to control plots. FAP plots also had significantly higher value yields (when including the value of all MHEP crops as well as main crops), and there was a significant correlation between pollinator abundance and yields (as measured by value of yields across all crops), and total income.

The differences in pollinator abundance between Gokwe South and Gokwe North districts seen in cycle 3 may be explained by regional differences in water availability. Despite efforts to select plots with similar biogeochemical environmental conditions, future research will need to ensure additional environmental variables are recorded to ensure comparability between plots.

The higher pollinator diversity and abundance in FAP plots gives an encouraging sign that FAP plots are beneficial to biodiversity as well as to the ecosystem services that those pollinators provide. Additionally, although impacts on non-pollinator diversity were not measured, anecdotal reports of benefits to fruit trees surrounding FAP fields were recorded.

As with many pilot studies, there are some points to improve on for future projects. Most significant of these will be ensuring that all farmers follow the plot design and farming techniques: lack of adherence to plot standard design and common harmonised farming practices by community groups resulted in almost half of the experimental plots being excluded from the data analysis. Despite this, the results of our research corroborate the findings of previous FAP research conducted by ICARDA. The visible effects of pollinator-friendly farming in the scope of the FAP trials can serve as motivation for farmers to reduce harmful behaviours that lead to pollinator decline, and indeed participating farmers were very positive about the FAP trial (Annexe 3). Such projects are not viable without farmer adhesion, so this is also an important outcome. More work is needed to build on the findings from this research pilot to complement and enhance evidence for FAP impact to support scaling and replication of the approach and pollinator-friendly farming.

This research has contributed to the on-going development of new agricultural guidelines by AgriTex and the Zimbabwe new national agroecology policy. It also supports international conventions and frameworks on biodiversity and agroecology, particularly as relating to biodiversity, diversification of income, input reduction, etc. However, for FAP to have the necessary space to scale up, it needs to be promoted through advocacy based on evidence that emphasize its value and potential in enhancing pollinators and increasing small-holder farmers’ yield and income. The FAP advocacy perspective aims to influence public policy decision-makers and donors, so FAP and other nature-based solutions are put back at the heart of national agricultural strategy and policies.

## 6. Acknowledgements

We are very grateful to the NAZ, ACF, Environment Africa and Agritex teams in the field.

## 7. Author contribution statement

The authors contributed to this paper in the following ways:

- S. Allebone-Webb: Data curation, formal analysis, visualisation, writing (original draft and reviewing/editing)
- F. Gossrau: Conceptualization, data curation, formal analysis, funding acquisition, investigation, Project administration, methodology, supervision, writing (original draft and reviewing/editing)
- C. Orland: Data curation, formal analysis, methodology, supervision, validation, writing (reviewing/editing)
- G. Bara: Conceptualisation, investigation, methodology, supervision, validation, writing (reviewing/editing)
- C. Fioekou: Project administration, supervision, validation
- A. Matsika: Conceptualisation, funding acquisition, investigation, project administration, resources, supervision, validation, writing (reviewing/editing)
- K. Riber: Funding acquisition, project administration, resources, validation
- B. Mahaman Dioula: Conceptualisation, Investigation, methodology, supervision, validation, writing (reviewing/editing)

## 9. Data availability

Anonymised data is on the mango platform https://zenodo.org/records/14802821 and is available on at.

## Annexe 1. Pollinator monitoring procedure and data collection

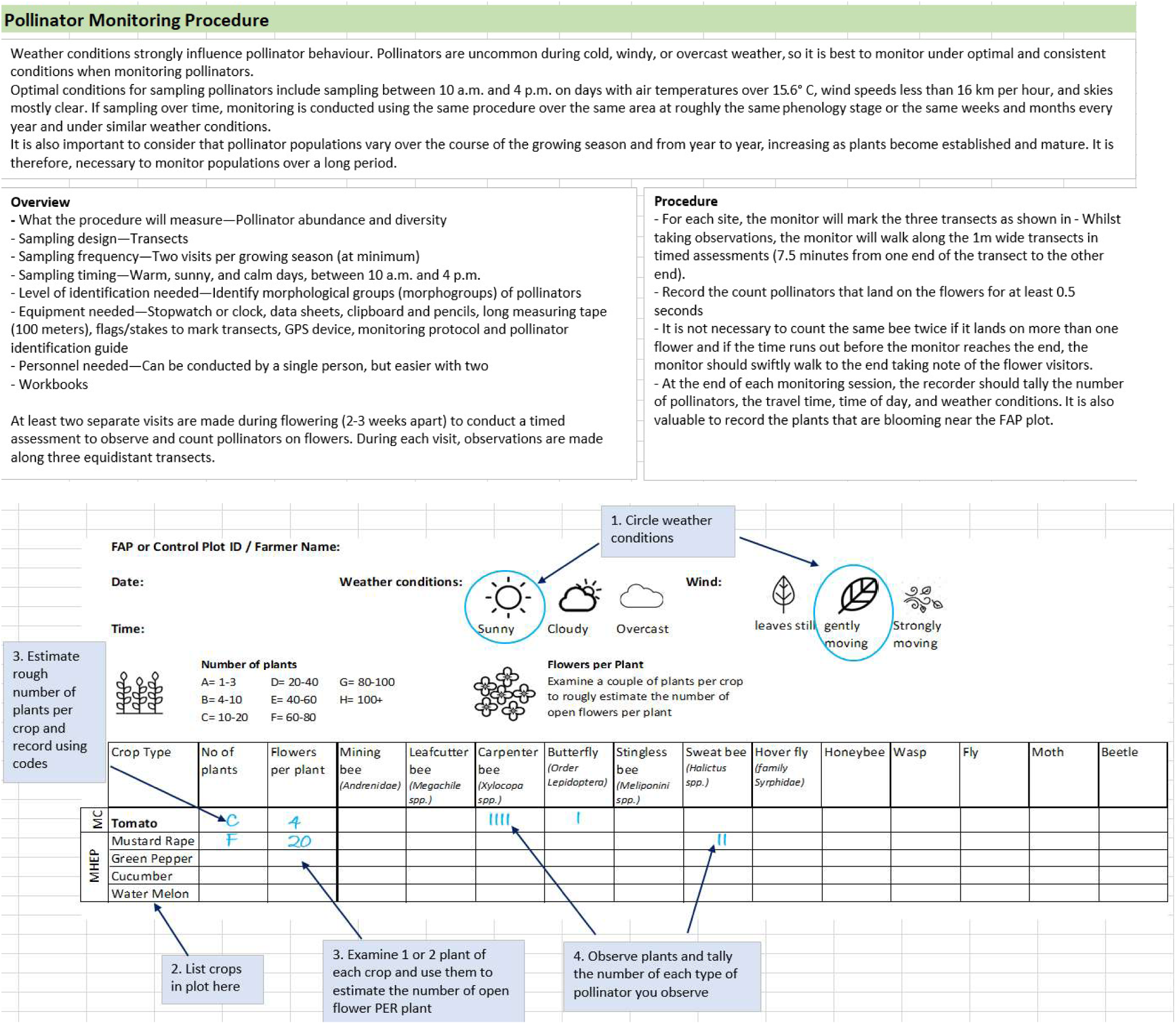

## Annexe 2. Additional results

**Table 2.** Analysis results of impact of treatment on pollinators by cycle (n = number of plots). Wilcoxon unpaired test in R.

|  |  | Pollinator abundance |  |  |  | Taxonomic richness |  |  | Pollinator diversity |  |  |
| --- | --- | --- | --- | --- | --- | --- | --- | --- | --- | --- | --- |
|  |  | n | median | IQR | W | median | IQR | W | median | IQR | W |
| Cycle 1 | FAP | 27 | 241 | 41 | W=0 | 12 | 0 | W=21.5<br>p<0.001 | 2.35 | 0.0751 | W=48<br>p<0.001 |
|  | Control | 14 | 29 | 7.25 | P<0.001<br>(p=2.17e-07) | 11 | 1 |  | 2.20 | 0.0925 |  |
| Cycle 2 | FAP | 28 | 154 | 32.5 | W=0<br>p<0.001 | 12 | 0 | W=84<br>P<0.001 | 2.41 | 0.0484 | W=36<br>p<0.001 |
|  | Control | 13 | 21 | 1 | (p=3.57e-07) | 11 | 1 |  | 2.34 | 0.0601 | (p=4.57e-05) |
| Cycle 3 | FAP | 24 | 133 | 47.5 | W=0<br>P<0.001 | 12 | 0 | W=38.5<br>P<0.001 | 2.39 | 0.152 | W=9<br>P<0.001 |
|  | Control | 11 | 18 | 25 | (p-value = 2.959e-06) | 10 | 2.5 |  | 2.11 | 0.109 | (p=1.344e-05) |

## Annexe 3. Acceptability of the FAP method

What is the likelihood for you to adopt FAP?

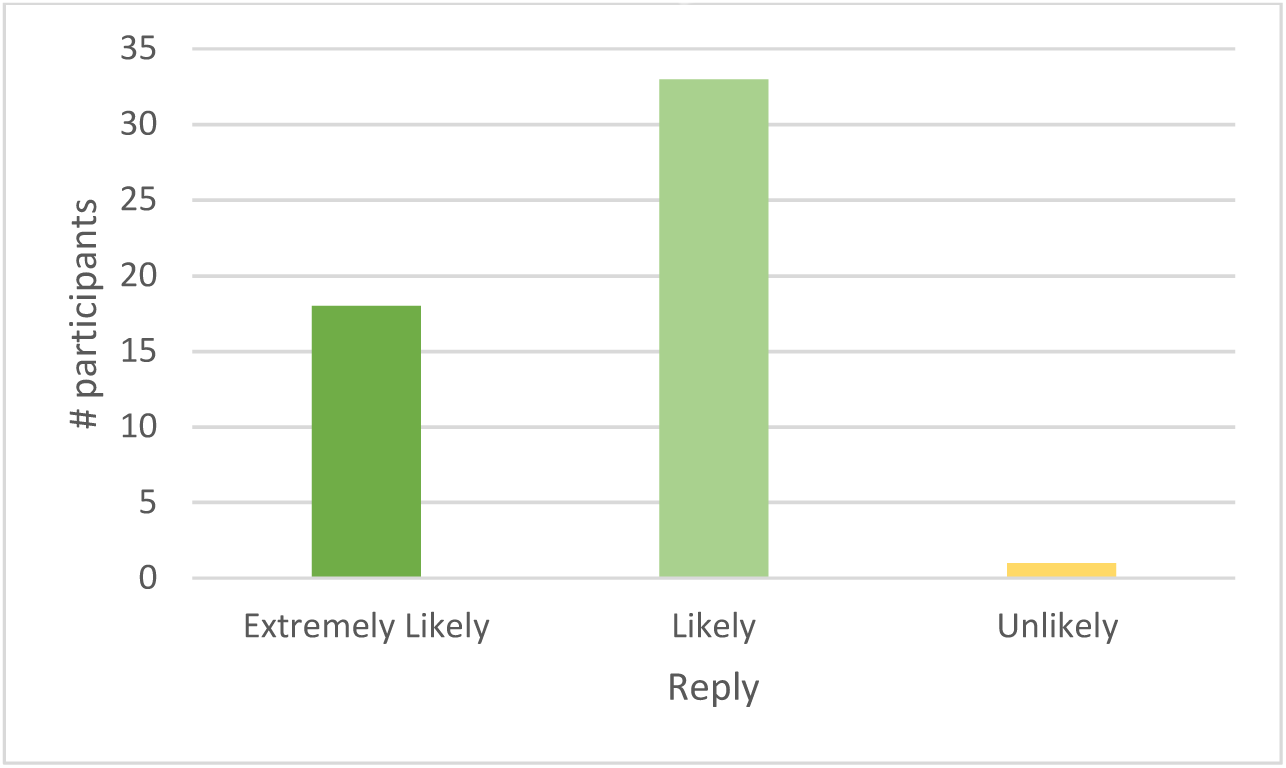

Is the cost of implementing FAP an issue that you would consider when implementing FAP?

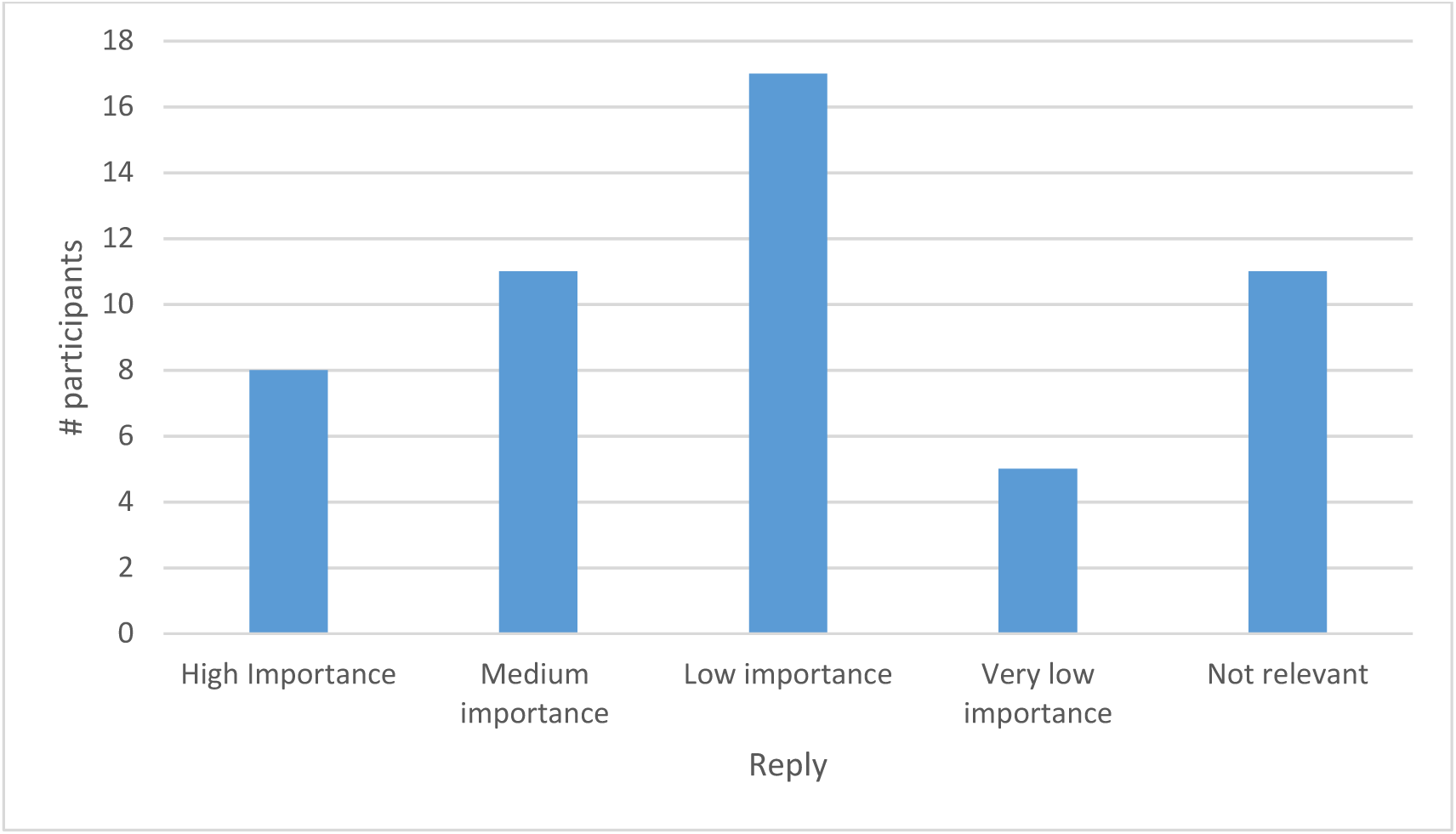

FAP is an easy technology, I can do it on my own

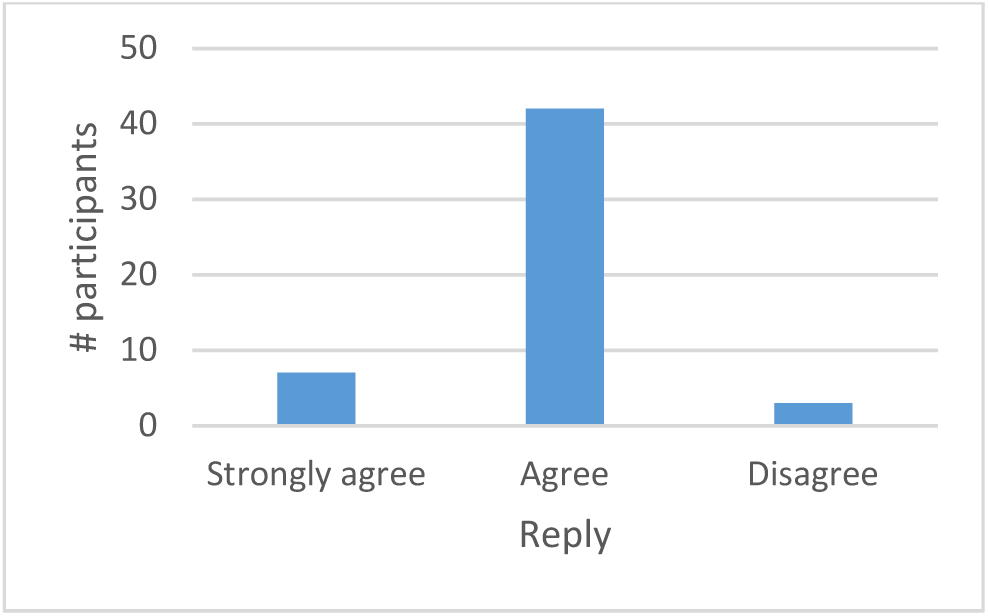

Everyone can do FAP

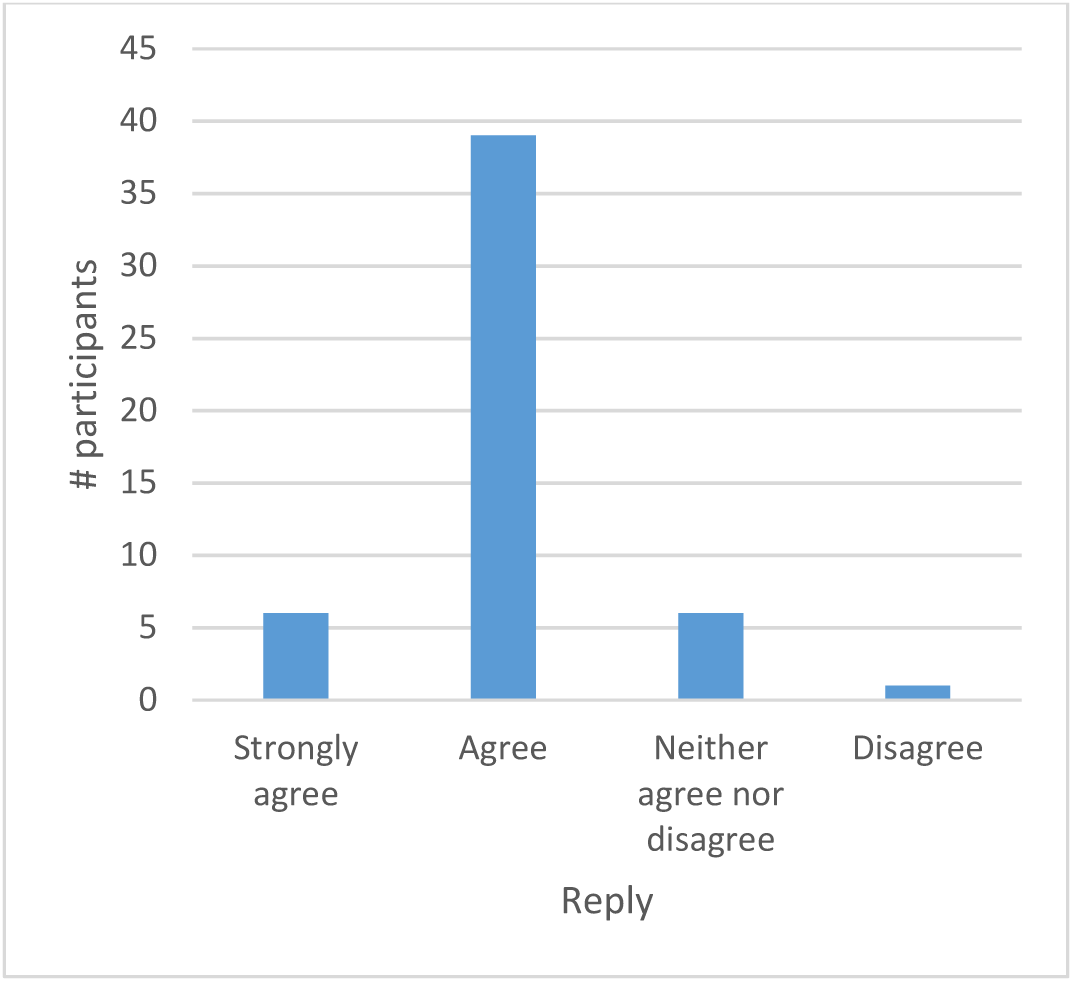

FAP is profitable

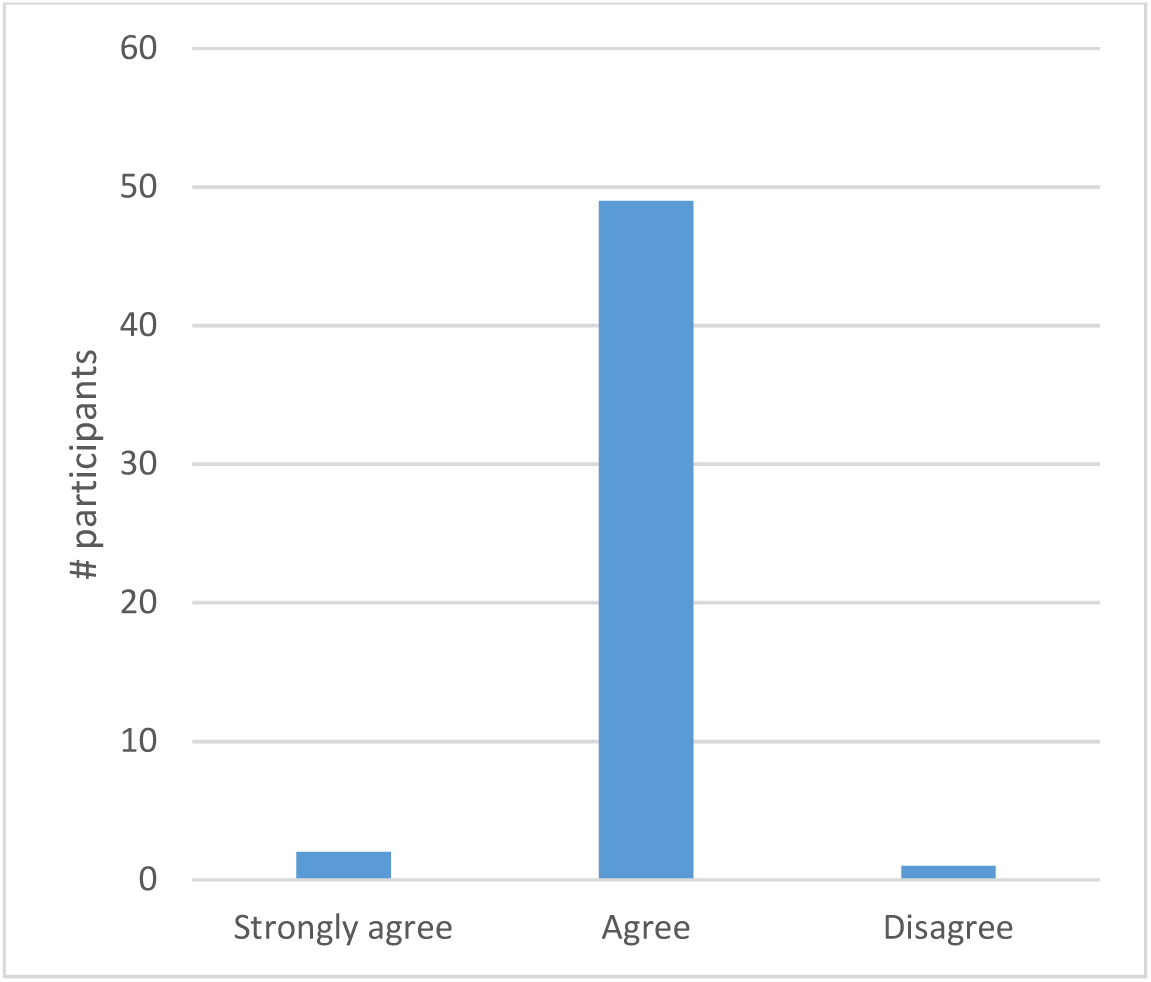

The conventional method of farming is more profitable than FAP

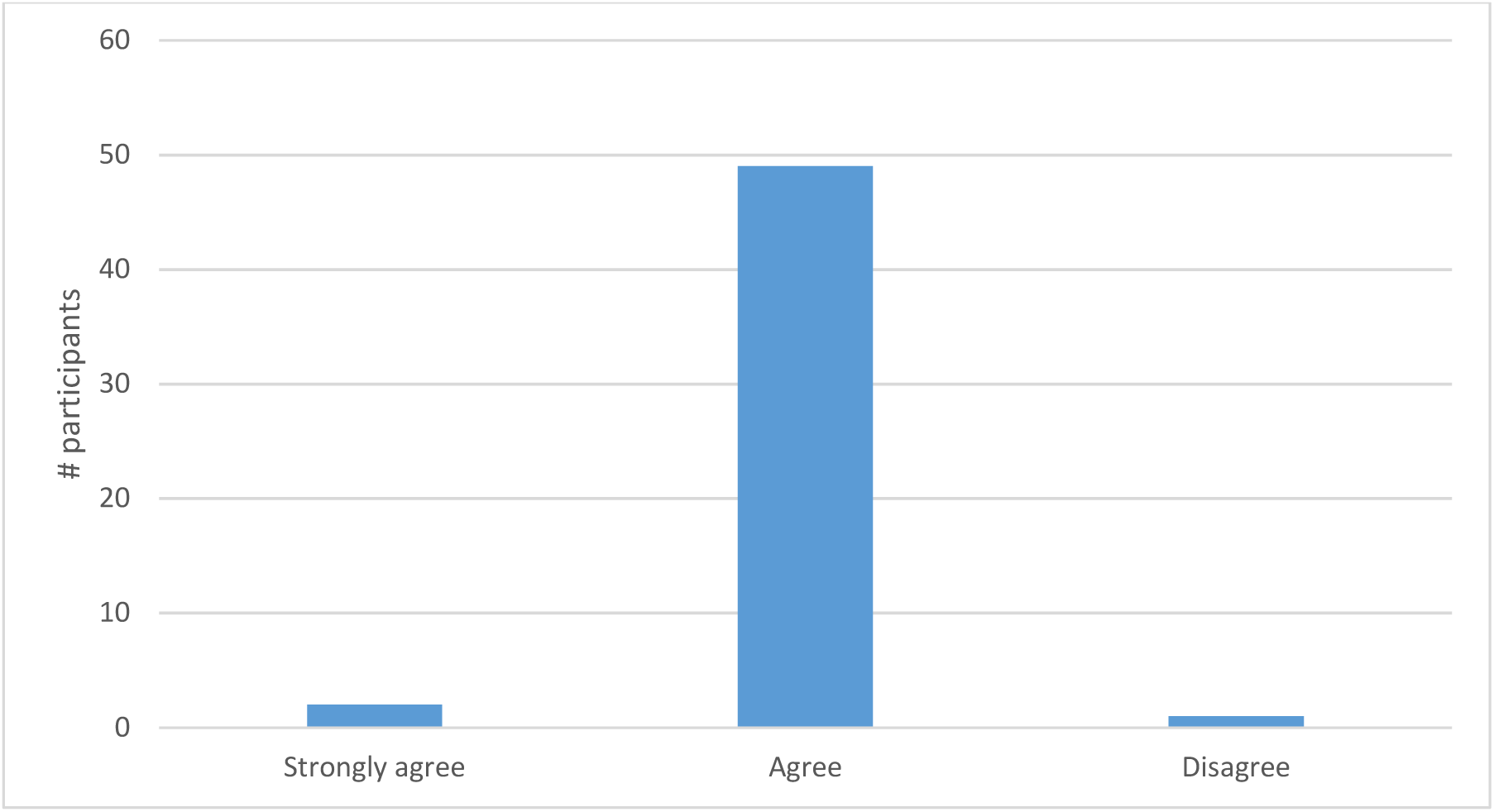

FAP is compatible with the land tenure systems in the area

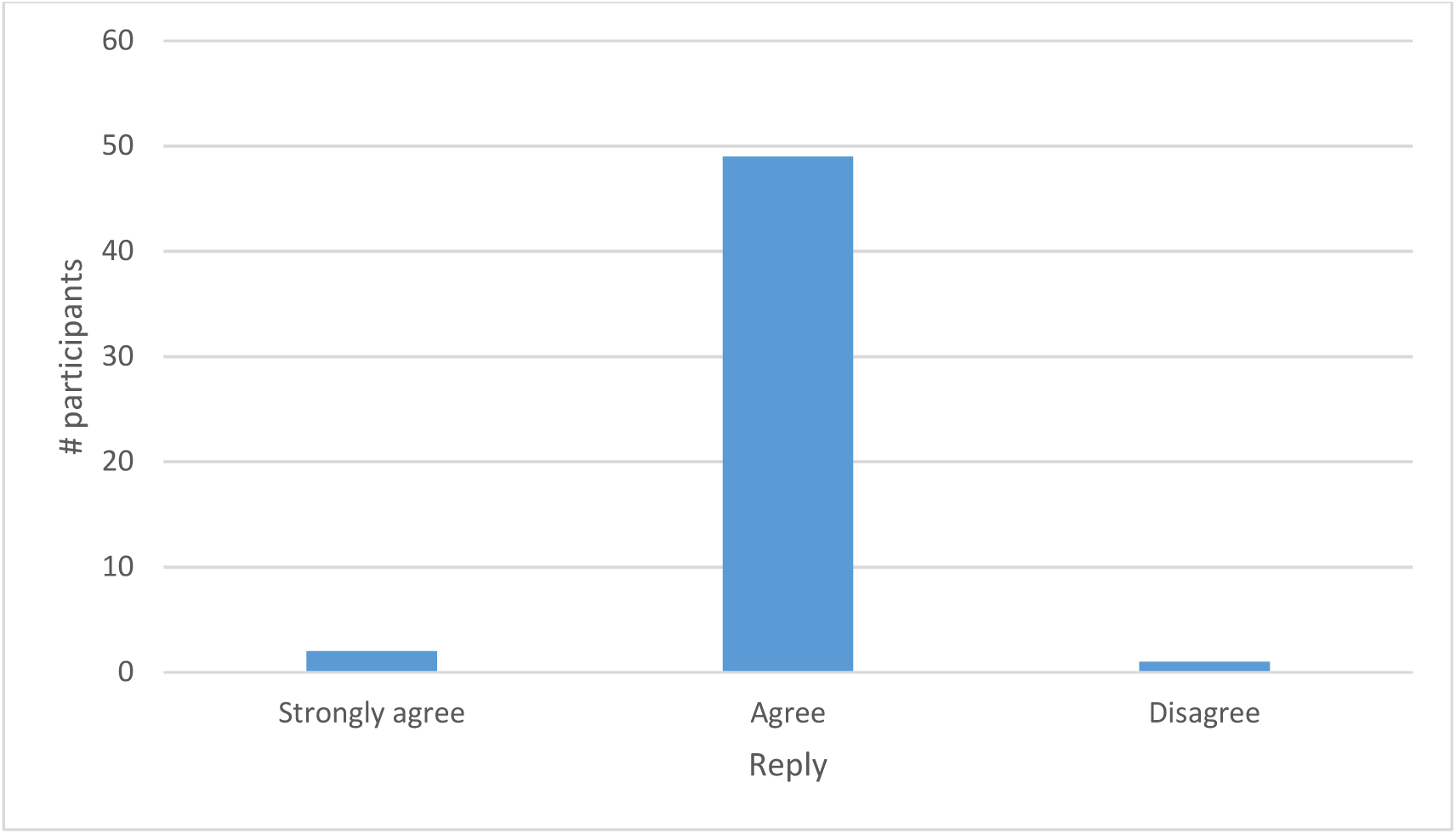

## Footnotes

4 an international aide NGO with the mandate to fight against hunger and its causes

5 “Principle 2: ACF research is ethically justified and scientifically valid” which includes “ACF obtains the informed consent of participants”, p.23.

6 Section 3.2.ii. “Consent”

7 P.1. 2) “Ensure that the consent is given of the persons concerned for each data collection”

